# Quorum sensing in Mycobacteria: understanding the recognition machinery conundrum through an *in-silico* approach

**DOI:** 10.1101/2024.03.06.583649

**Authors:** vani Janakiraman, Krovvidi Phani Sarath Teja

## Abstract

Bacteria employ a cell-to-cell communication process called quorum sensing (QS) to orchestrate group behaviors like exo-factors and host-adapted traits. The QS machinery in gram negative bacteria comprises of LuxR proteins (and their homologs) that are transcription factors which recognize and bind to the classical signaling molecules acyl homoserine lactones (AHLs). On the other hand, QS in gram positive bacteria is mediated through autoinducer peptides recognized by two-component systems (TCS). However, in acid-fast bacteria, the very process of QS and the underlying molecular machinery remains elusive.

In the present work, we have investigated the proteins annotated as LuxR family proteins of the clinically important genera of the acid-fast bacteria, mycobacteria through computational analysis. We have chosen *Mycobacterium tuberculosis* (Mtb), the etiological agent of tuberculosis and a most widely used model system for Mycobacterial studies, *Mycobacterium smegmatis*. A total of 17 genes annotated as LuxR homologs (7 from Mtb and 10 from *M. smegmatis*) were analyzed. We found that only 14 of these proteins (5 from Mtb and 9 from *M. smegmatis*) harbor the HTH motif typical to the LuxR/FixJ superfamily of transcriptional regulators affirming their belonging to LuxR family. Rv0894 and MSMEG_0545 both annotated as LuxR homologs, do not harbor HTH motif and RegX (also annotated as LuxR homolog) does not have the tetra helical HTH which is the characteristic of LuxR/FixJ superfamily and hence are not LuxR family proteins. Interestingly, most of the LuxR family proteins (2 in Mtb and 6 in *M. smegmatis*) are response regulators (RRs) that harbor REC domain that is involved in phosphotransfer from the histidine kinases (HK) thus forming a TCS involved in physiological processes. Few of them have their cognate HKs while few are orphan regulators. The remaining of the proteins harbor various sensory domains that include MalT, PAS, GAF, AcyC, ATPase, TPR, TOMM, and HchA which are either enzymes or bind to small ligand or proteins. STITCH-an online protein-chemical interaction server in deed revealed various small molecules including c-di-GMP (QS molecule in *M. smegmatis*), and 3-oxo-C12-HSL (a QS signal in *P. aeruginosa*) binding to the ligand-harboring LuxR proteins. Our study not only confirms the authenticity of Mycobacterial LuxRs but also reveals the diversity of domains in the proteins annotated as LuxR family members in mycobacteria. This type of domain organization is strikingly different from the classical quorum sensing machinery of other bacteria, which might have evolved for a hitherto unknown multifunctionality including QS.

**IMPORTANCE:** Though QS is an important biological process regulating various traits in most other bacteria, the workings of it remain elusive in Mycobacteria. Hence, in the present study, we have attempted to unearth the nature of proteins annotated as LuxR family proteins (which participate in quorum sensing in other bacteria) in mycobacteria through *in silico* analysis. We show that LuxRs of mycobacteria fall into four different families of LuxR/FixJ group of proteins, based on the presence and nature of the sensory domains. Our results provide an understanding of how diverse LuxR proteins could be in terms of domain composition and hence function. This also hints towards the ligands of varied nature such as second messengers and aromatic compounds that might potentially bind to some of these LuxRs harboring the GAF/PAS domains and thus participate in QS or in stress-response phenomena suggesting that these mycobacterial proteins might have in other physiological processes important for survival of the bacteria as an individual or as a community in various.

## INTRODUCTION

Bacterial cell-to-cell communication classically known as quorum sensing (QS) is a cell density dependent phenomenon wherein bacteria detect changes in population density via chemical messengers and respond collectively to perform various physiological functions. These messengers called autoinducers accumulate in the surrounding milieu and upon reaching a certain threshold, trigger transcriptional regulation of genes which usually encode for products that serve as public goods^1^. Typically, traits like bioluminescence^2^, biofilm formation^3^, exo-factor production^4,5^, competence^6^, sporulation, conjugation^7,8,9^, and swarming^10^ amongst others are well established QS regulated phenotypes. Exhibiting these traits is usually energy consuming and thereby expensive and unproductive when performed by a single cell but are efficient when performed as a group^11^. While gram negative and gram-positive bacteria have different types of machinery for synthesizing, secreting, and detecting these autoinducers, the response is always group behavioral. QS provides an advantage to the bacteria living with their hosts or living as polymicrobial communities and allows bacteria to function as multi-cellular organisms^12^.

Gram negative bacteria utilize acylated homoserine lactones (AHL) as signaling molecules^13^ and the Lux system for synthesizing and detecting the AHLs^14^. The Lux system was named after the Lux operon of *V. fischeri* where QS was first identified^2^. AHLs are known to be synthesized by LuxI synthases and its homologs which then diffuse freely out of the cells. Upon reaching a certain concentration threshold, they are rapidly internalized and bind to transcription factors called LuxR and its homologs. The LuxRs are highly specific to particular AHLs and only get activated upon binding to their cognate AHLs^15^. Gram positive bacteria on the other hand employ modified peptides^16,17^ as signals which bind to sensory receptor proteins on the membrane that are a part of two-component system (TCS). The receptors are histidine kinases (HK) which upon ligand binding trigger the phosphorylation of the conserved histidine residue^18^. The phosphate is then transferred to a response regulator (RR) (which is the second component of the TCS) also a transcription factor, thus getting activated^19^. Upon activation, the RRs regulate the transcription of genes under their control culminating in exhibiting QS traits^20^. While AHLs and peptides are the most prevalent QS signaling molecules among bacteria, various other secreted molecules were shown to be acting as autoinducers^21,22,23,24,25,26,27,28,29^. The most prominent ones are auotinducer-2 (AI-2) and γ-butyrolactones (GBLs). AI-2 is the common language for both gram positive and gram negative bacteria and mediates communication among different species of bacteria^21,30,31^. GBLs on the other hand are specific to the phylum *Actinobacteria* that controls the production of various antibiotics^29^. Surprisingly, many members of *Actinobacteria* also harbor proteins that are annotated as LuxRs^32^ that typically are the recognition systems for AHLs in gram negative bacteria. Neither their nature nor the function is ever visited.

LuxR proteins are a part of LuxR/FixJ superfamily (referred as LuxRs from here-onwards) of transcriptional regulation proteins and are typified by the presence of a signal receiver domain in the N-terminus and a DNA-binding helix-turn-helix (HTH) domain in the C-terminus^33^. This superfamily of proteins is divided into various subfamilies based upon their mode of activation^34^. One of the classes is the classical LuxRs of gram negative bacteria harboring AHL binding domain which upon binding to AHLs get activated and enhance transcription of downstream genes and in rare cases get repressed^35^. Usually, LuxR genes are located adjacent to the LuxI genes forming a cognate pair. Interestingly, few LuxR genes exist as solos with no cognate LuxI found in the genome^36,37^. These have evolved to recognize AHLs from other bacteria or molecules other than AHLs. With so far available reports, Actinobacterial LuxRs seem to be solo LuxR genes with no LuxI genes^32^. *Mycobacterium tuberculosis* (Mtb) which is a clinically important member of this phylum responsible for causing tuberculosis harbors as high as 7 different genes annotated as LuxRs^38^ with no known cognate LuxIs and so is its saprophytic counterpart *Mycobacterium smegmatis* which harbors 10 genes annotated as LuxRs. Interestingly, in addition to the annotated LuxRs, *M. tuberculosis* harbors 3 more proteins and *M. smegmatis* has 24 more proteins though not annotated as LuxRs but possess similar domain architecture as that of annotated LuxRs^32^. Intriguingly, all of the mycobacterial LuxRs are not the classical LuxR proteins like the ones in gram negative bacteria and harbor other unique domains. The presence of these domains and absence of AHL binding domain posits a possible different mode of activation of these LuxRs by ligands of different chemical nature and their role in several other physiological processes including QS. Interestingly, there is an indirect evidence of QS in mycobacteria where second messengers have been shown to mediate biofilm formation^39^ for which the first messengers (extracellular signaling molecules) are unknown and *whiB3*-a putative transcriptional regulator whose levels were shown to be inversely correlating with bacterial density thus might be a QS regulated molecule^40^, the signaling molecules and their recognition systems mediating QS are still unknown in mycobacteria. In this study, we have attempted to tease out the nature of QS machinery in the clinically important member of *Actinobacteria*-Mtb and its non-pathogenic equivalent *M. smegmatis* using a computational approach.

## RESULTS

LuxR proteins form the quorum sensing machinery in large number of bacterial genera. Mtb and *M. smegmatis* harbor proteins have only been annotated as LuxR family and there are no experimental evidences to prove their functionality. In the present work, mycobacterial LuxR proteins have been computationally analyzed for the presence of signal binding domains and DNA binding domains which is the signature of classical LuxRs. We have tried to delineate the true nature and versatility of these domains and plausible ligands they might bind to in order to gain insights into the process of quorum sensing in mycobacteria.

### Mycobacteria harbor unusually high number “LuxR family” genes

A total of 7 proteins in Mtb and 10 proteins in *M. smegmatis* have been annotated as LuxR family proteins or possibly LuxR proteins. Strikingly, these proteins range from 85 amino acids to 1137 amino acids in length whereas most of classical LuxR proteins/LuxR homologs range from 230 to 270 amino acids. The domains these proteins harbor is illustrated in figure 1. Majority of proteins, harbor CitB superfamily that contains REC and HTH domains making REC the most commonly associated domain with HTH (since all typical LuxRs are transcription factors and hence harbor HTH) among LuxRs. Other domains present in these proteins include REC, MalT, PAS, GAF, AcyC, and ATPase. Detailed analysis of these domains and their plausible functions are discussed in other sections of the results.

**FIG 1.**
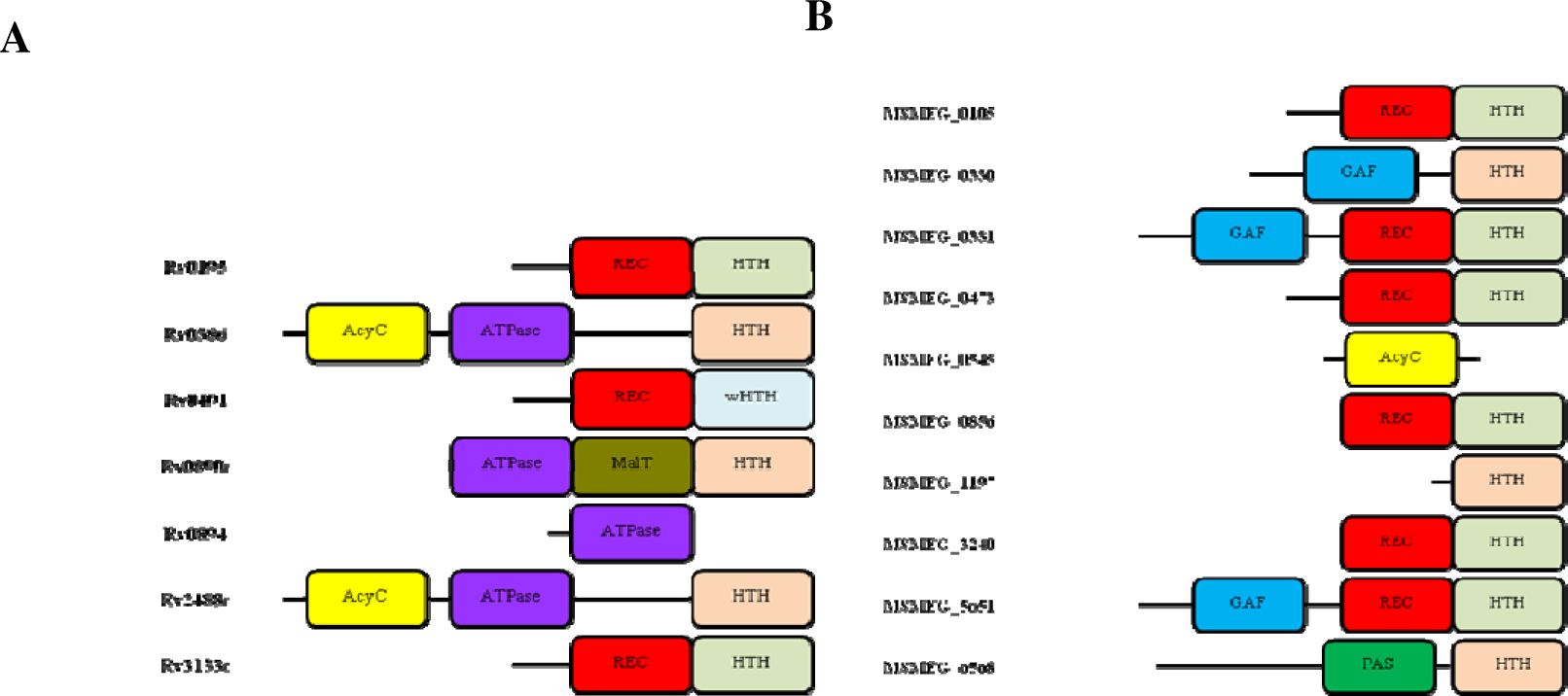
Domain architecture of mycobacterial LuxR proteins from (A) *M. tuberculosi*s and (B) *M. smegmatis*. Representations of the domains and so the protein lengths are not to the scale. **HTH**-DNA binding helix-turn-helix motif, **wHTH**-winged helix-turn-helix motif, **REC** (receiver) or CheY-like phosphoacceptor-signal receiver domain, **AcyC**-class 3 adenylate cyclase domain, **ATPase**-ATPase domain, **MalT**-maltotriose, and ATP binding domain, **GAF**-ligand binding domain named after c**G**MP-specific phosphodiesterases, **a**denylyl cyclases and **F**hlA proteins that harbor this domain, **PAS**- **P**er- **A**rnt-**S**im, a sensory domain for protein-protein or protein-small molecule interactions. It may be noted that mycobacterial LuxRs also harbor atypical domains (AcyC, GAF, PAS, and ATPase) in addition to the classical domains (REC, and MalT) in LuxR/FixJ superfamily of transcription regulators.

To know if the LuxRs in Mtb and *M. smegmatis* are orthologs of one another, the same was searched in Mycobrowser (data not shown). Rv0491 of Mtb has an ortholog in *M. smegmatis* which is MSMEG_0937 (regX3) which is not in the annotated list of LuxR family. Rv3133c on the other hand has two orthologs in *M. smegmatis* which are MSMEG_3944, MSMEG_5244 (both annotated as devR) neither of which have been annotated as LuxR family protein. MSMEG_0105 has an ortholog in Mtb which is Rv0844c and codes for narL and is also not been annotated as LuxR family protein. It appears that all of the LuxRs are not conserved among the pathogenic and non-pathogenic members of mycobacteria.

### Mycobacterial LuxR family proteins harbor HTH that are characteristic of LuxR/FixJ superfamily of transcription regulators

The characteristic of LuxR/FixJ family of proteins is the HTH domain and most of the mycobacterial LuxRs harbor HTH which usually is in association with sensory domains. Majority of these HTH are associated with REC domain forming CitB superfamily of proteins (CDD ID Cl28474) indicated as light green boxes in figure 1. The remainder of the proteins has HTH that are the sub class of HTH superfamily (CDD ID Cl21459) indicated as light red boxes in figure 1. Irrespective of the different classifications, both of them belong to LuxR/FixJ superfamily of transcription regulators. The sequence alignment for other well characterized members of LuxR/FixJ superfamily was performed (data not shown) and the HTH region of the mycobacterial LuxRs is aligned with one representative member of LuxR/FixJ superfamily of transcription factors (LuxR of *A. fischeri*) to check for the signature motifs. Most of the LuxRs of mycobacteria have consensus amino acids (indicated by arrows) in the HTH region as shown in fig 2. In addition, secondary structures of these regions revealed the presence of 4 alpha-helices which is the characteristic of LuxR/FixJ superfamily of transcription factors (figure 2). A further analysis of the HTH region revealed that 3 proteins do not harbor the classical LuxR/FixJ type HTH. RegX3 (Rv0491) which is one of those proteins, has winged HTH. It is a part of OmpR superfamily that contains 3 alpha helices and a wing. The sequence alignment of Rv0491with OmpR superfamily of proteins OmpR and PhoB is shown in fig 2E. Hence, Rv0491 which does not harbor the characteristic four helix of LuxR/FixJ superfamily is not a LuxR family protein. Two proteins Rv0894 and MSMEG_0545 do not harbor the HTH domain and only have ATPase and AcyC domains respectively hence making them non LuxR family proteins. All together, out of 17 proteins analyzed, only 14 are true LuxR family proteins whereas RegX3, Rv0894 and MSMEG_0545 do not belong to this family and hence were not considered for further study.

**FIG 2.**
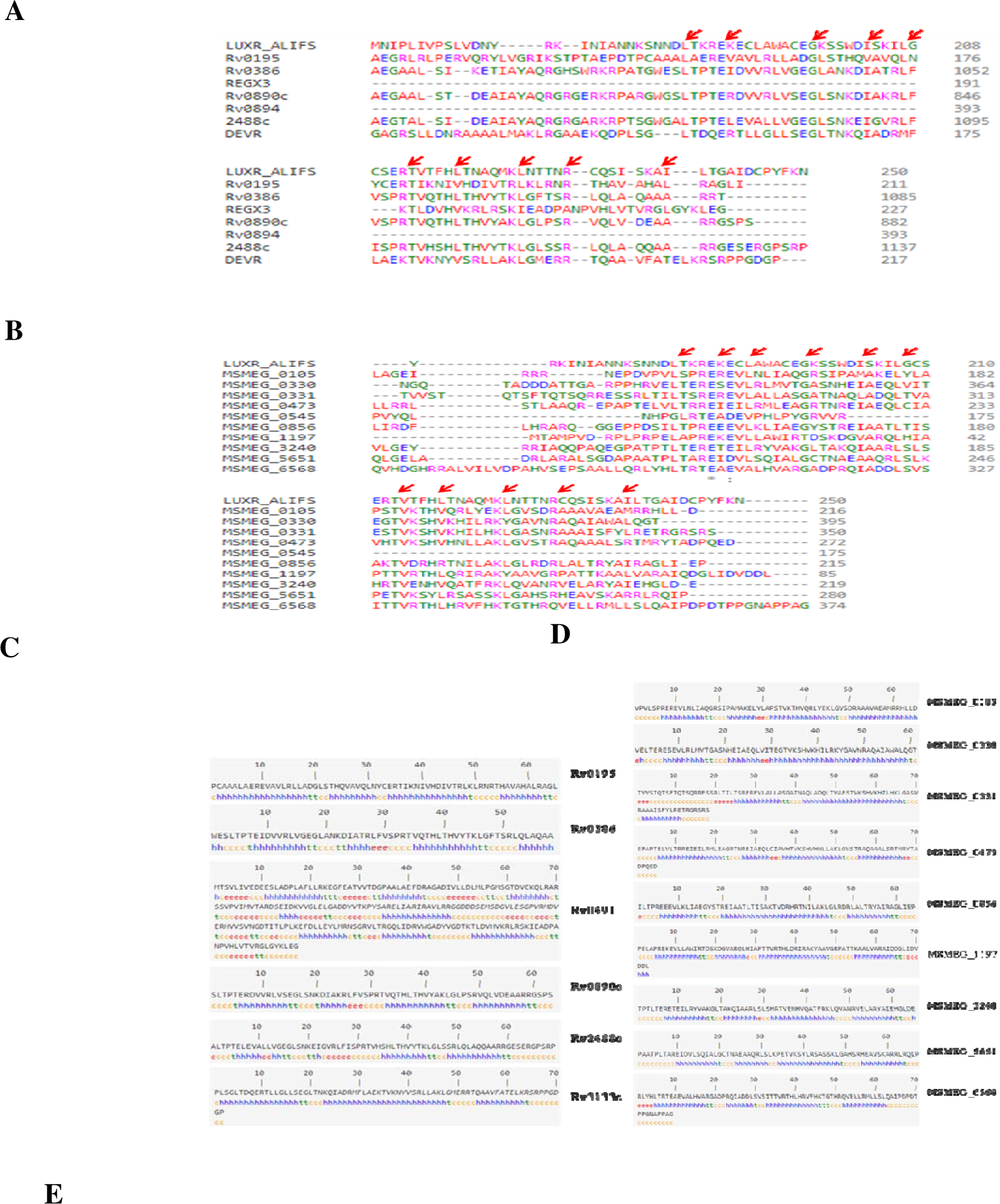
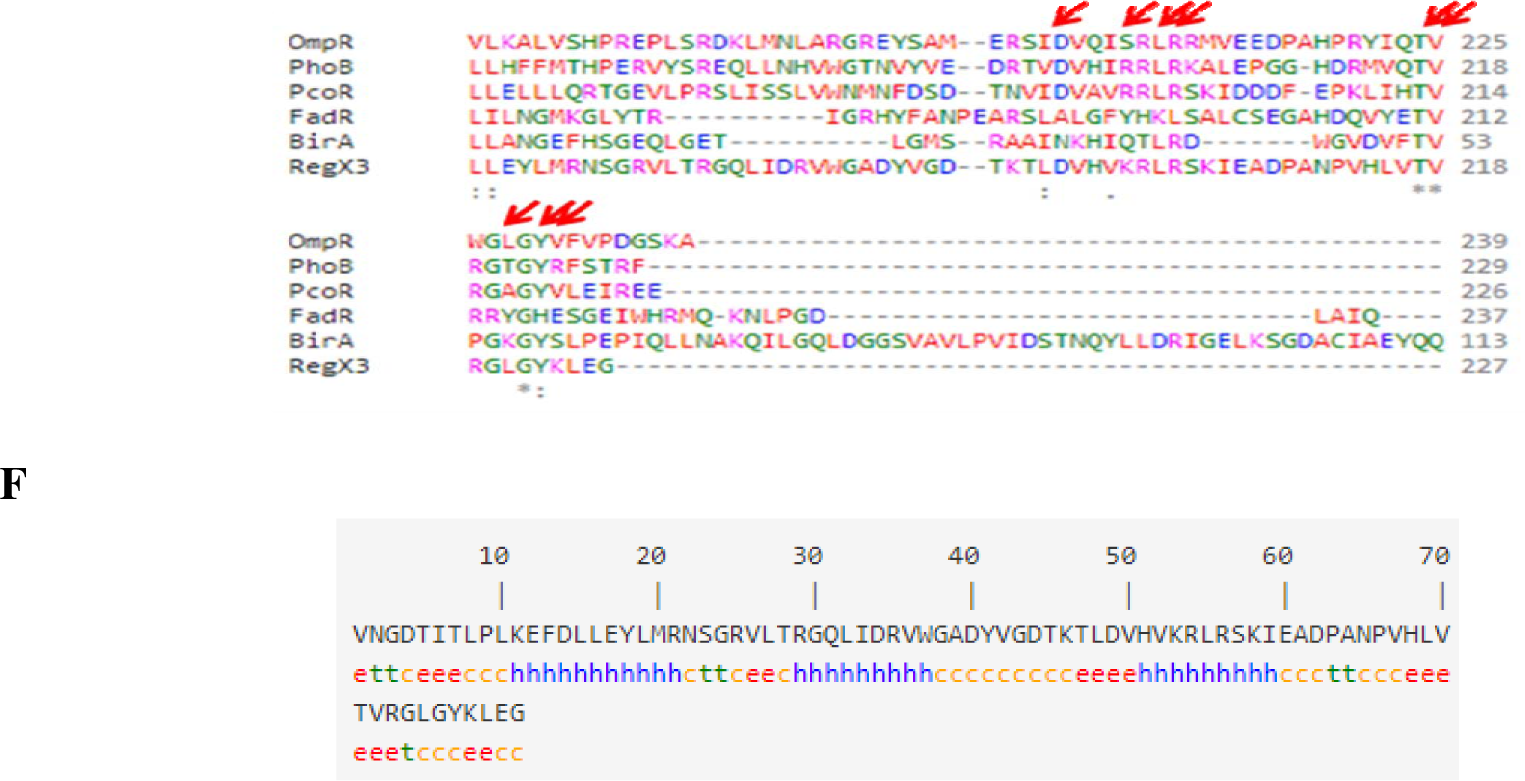
Sequence alignment and secondary structure prediction of HTH region of LuxR proteins in mycobacteria. Consensus amino acids are indicated by arrows. (A) Sequence alignment of Mtb LuxRs with the classical LuxR (a well characterized LuxR family protein of *A. fischeri* used as a representative for LuxR/FixJ superfamily). Only 5 of 7 LuxRs (Rv0195, Rv0386, Rv0890c, 2488c and devR) have conserved amino acid as that of LiuxR from *V. fischeri.* (B) Sequence alignment of 10 LuxR family proteins of *M. smegmatis* with LuxR. Only 9 out of 10 proteins in *M. smegmatis* have the consensus amino acids in the HTH region that are characteristic of LuxR/FixJ superfamily of proteins. The three proteins (Rv0491, Rv0894 of *M. tuberculosis* and MSMEG_0545 of *M. smegmatis*) lack the consensus amino acids and hence might not belong be LuxR/FixJ superfamily of transcription regulators Sequences of LuxR family proteins of (C) *M. tuberculosis* and (D) *M. smegmatis* showing the typical four helices of HTH motif (indicated as blue text) as analyzed by secondary structure prediction tool-SOPMA. (E) Sequence alignment for HTH domain of RegX3 with well-characterized winged-HTH (wHTH)-harboring proteins. RegX3 Harbors the consensus amino acids of the wHTH-containing proteins hence making it not a member of LuxR/FixJ superfamily of transcription regulators. OmpR, PhoB, PcoR, FadR, and BirA are all transcription factors from *E. coli* whose C-terminus was considered except for BirA which harbors HTH at N-terminus. (F) Predicted secondary structure of C-terminus of RegX3 showing the typical 3 helices of WHTH and a coil (indicated as red text) at the end of the protein. This analysis confirms that RegX3 indeed belongs to OmpR family of HTH but not to LuxR family.

### LuxRs of mycobacteria do not contain AHL binding domain but harbor other sensory domains typical of LuxR/FixJ superfamily

Two major families of LuxR/FixJ superfamily of proteins are the AHL-binding domain and REC domain containing proteins. None of the Mycobacterial LuxR harbor AHL-binding domain. But, majority of the mycobacterial LuxRs harbor REC domain. REC (receiver) or CheY-like phosphoacceptor in association with HTH is a common feature of bacterial two component systems. In this context, it is interesting to note that 8 of the mycobacterial LuxRs (Rv0195, Rv3133c, MSMEG_0105, MSMEG_0331, MSMEG_0473, MSMEG_0856, MSMEG_3240, and MSMEG_5651) harbor REC domain in combination with HTH. However, few of them (MSMEG_0331, and MSMEG_5651) in addition to harboring REC and HTH also harbor GAF domain (a type of signal binding domain). The presence of two sensory domains might explain dual mode activation and hence function of these proteins. The functional significance of GAF domain is explained in latter section.

Of the eight REC-harboring proteins, Rv3133c (devR) of Mtb has been both structurally and functionally characterized and is a part of two component system with a cognate histidine kinase Rv3132c (devS). This kinase interacts with its cognate response regulator (Rv3133c) by transferring the phosphate from histidine residue (present in HK) to aspartate residue (D54) in Rv3133c thereby regulating transcription. The sequence alignment for other well characterized REC-harboring proteins was performed (data not shown) and the REC domain of the mycobacterial LuxRs was aligned with one representative member of REC-harboring proteins (CheY of *E. coli*) to check for the signature motifs. The sequence alignment for all REC-containing mycobacterial LuxRs is presented in figure 3 which revealed that Rv0195 of Mtb does not harbor the consensus amino acids (especially the aspartate) of the REC domain while DevR harbors the conserved aspartate (D54) that receives phosphate from DevS^41^. In *M. smegmatis*, MSMEG_0331, MSMEG_0473, and MSMEG_5651 do not harbor the consensus amino acids of REC domain while the other 4 proteins do harbor those amino acids (figure 3). Ironically, Rv0195 is a well recognized member of Mtb two-component systems. So, to validate our results and also since the REC domain harbors the aspartate residue that receive phosphate from histidine kinase, we hypothesize that these REC containing mycobacterial LuxR family proteins might be the response regulators (RR) of two-component system. A search for the TCS of mycobacteria using p2cs online browser, and from literature^42^ revealed the presence of 10 paired TCS and 8 orphan proteins in Mtb. *M. smegmatis* has as many as 23 paired TCS, 22 orphan proteins and 1 triad system (data not shown). The list contained the characterized TCS pair Rv3133c-Rv3122c (DevR-DevS) and Rv0491-Rv0490 (RegX3-SenX3) systems although RegX3 is not a member of LuxR. But, in support to our domain analysis for REC, Rv0195 is not a part of the p2cs TCS list in Mtb hence confirming that Rv0195 harbors a different domain but not REC. Surprisingly, the protein Rv0844c (identified in p2cs) a part of paired TCS in Mtb has not been annotated as LuxR family but has the same domain architecture as REC-harboring LuxR thus expanding the list of known LuxRs in Mtb.

**FIG 3.**
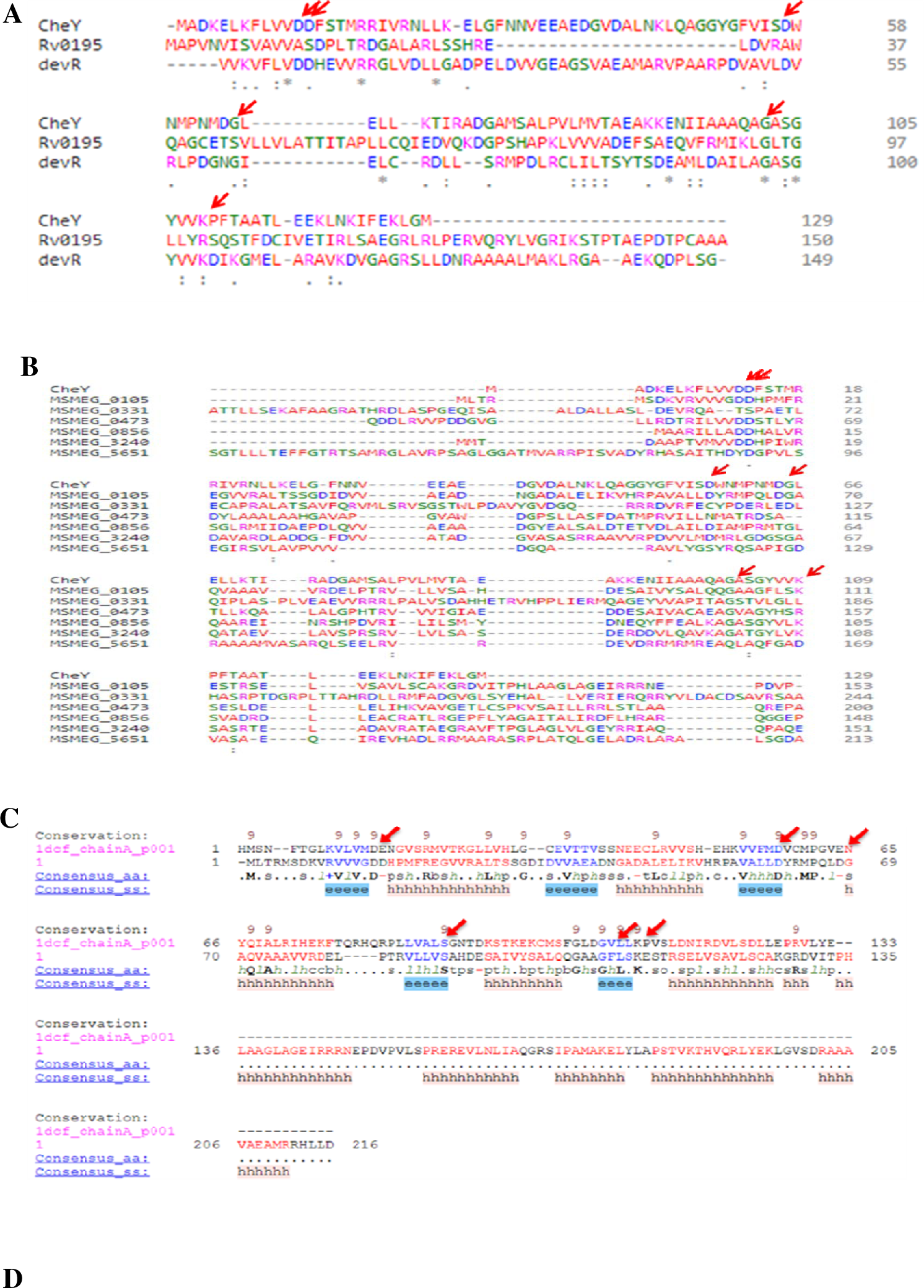
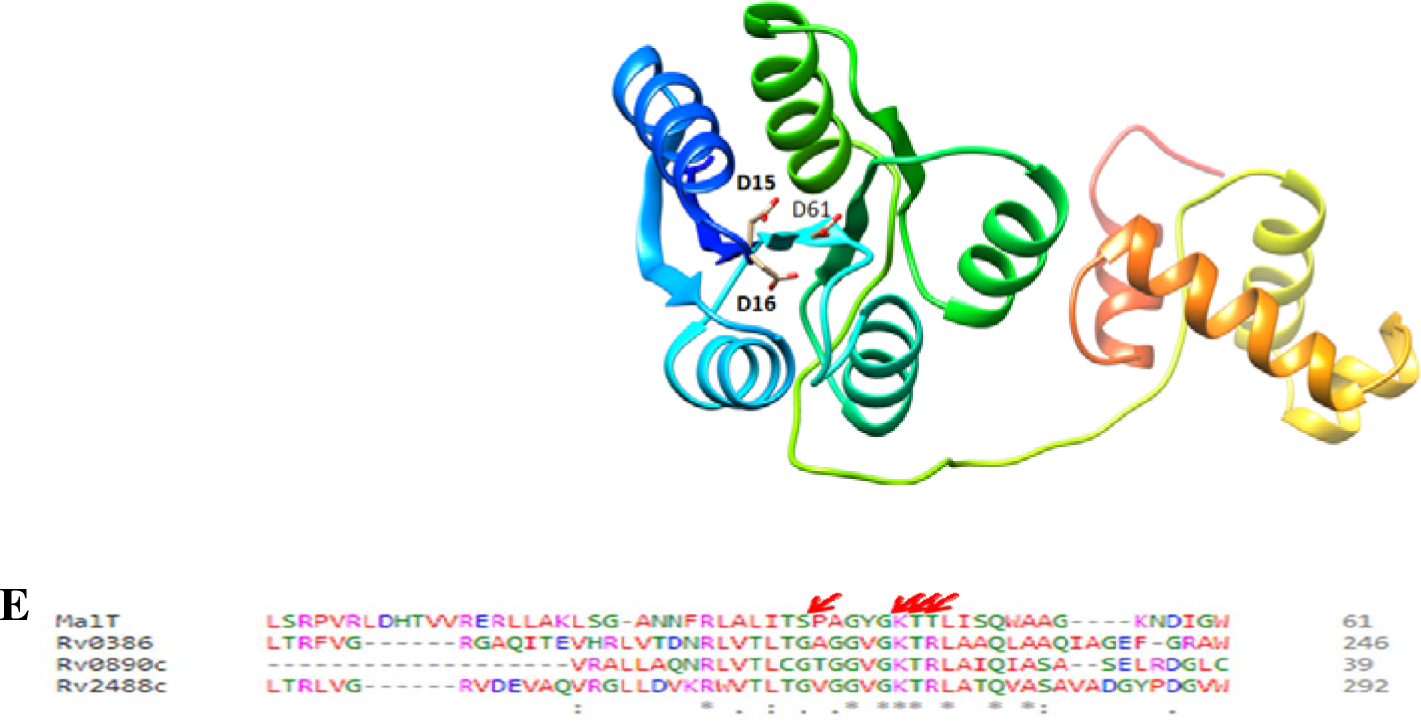
Sequence alignment and structure prediction of REC and MalT domain of LuxRs in mycobacteria. Consensus amino acids are indicated by arrows. (A) *M. tuberculosis* has 2 REC domain harboring LuxR family proteins and only 1 of them (devR) has conserved amino acids as that of CheY (a well characterized REC domain harboring protein of *E. coli* used as a representative). (B) Sequence alignment of 6 LuxR family proteins of *M. smegmatis* with REC domain. Out of 6 REC-containing proteins in *M. smegmatis*, MSMEG_0331 and MSMEG_5651 do not have the consensus amino acids (aspartate) in the REC domain needed for accepting the phosphate group from histidine kinase. Hence Rv0195, MSMEG_0331 and MSMEG_5651 though being LuxR proteins, do not harbor REC domain and hence might not be a part of two-component systems. (C) Structure-based sequence alignment of MSMEG_0195 (representative for all the REC-harboring LuxRs in mycobacteria) with REC-harboring protein of *A. thaliana*. (D) Predicted structure of the MSMEG_0105. The conserved aspartate residues at positions 15, 16 and 61 are the most probable sites for phosphotransfer from histidine kinases. (E) Multiple sequence alignment for MalT /ATPase domain harboring LuxRs (Rv0386, Rv0890c, and Rv2488c) with MalT of *E. coli* (a representative for LAL (large ATP binding regulators of the LuxRs). All the 3 proteins harbor a motif called Walker A motif (**G**XXXX**GKS/T**, where X is any residue, indicated as arrows) which is typical of LAL family. Hence all the three proteins make up one group of mycobacterial LuxRs which is LAL family.

*M. smegmatis* on the other hand has 4 of the annotated LuxRs as part of TCS. MSMEG_0105, MSMEG_0856 and MSMEG_3240 have cognate HK-MSMEG_0106, MSMEG_0854 and MSMEG_3239 respectively whereas; MSMEG_0473 (for which there were no consensus amino acids identified in REC domain) is an orphan TCS member with no known cognate HK. MSMEG_0331 and MSMEG_5651 are not included in the TCS list of *M. smegmatis* in p2cs database in agreement with our results of REC-domain sequence alignment. The neighboring genes for MSMEG_0473 do not code for HKs hence making it either a solo or not a REC-harboring protein at all. Similar to Mtb protein Rv0844c (identified in p2cs) which is not annotated as LuxR yet harbor LuxR type HTH, 5 proteins in *M. smegmatis* (MSMEG_0459, MSMEG_0983, MSMEG_1494, MSMEG_4969, and MSMEG_6236) also harbor LuxR type HTH. MSMEG_0983 is the only solo RR out of the 5 newly identified LuxRs but its gene neighborhood harbors MSMEG_0980 that codes for HK although MSMEG_0981 (adjacent gene to MSMEG_0980) codes for RR. This could be that the LuxR-MSMEG_0985 is a part of triad system comprising one HK and two RRs. The remaining of the LuxRs have their cognate HKs (data not shown). To further validate the results for the REC-harboring LuxRs, a sequence-structure alignment of MSMEG_0105 (representative for all REC-harboring LuxRs) with the structure representing the REC domain (PDB ID: 1DCF) of ethylene receptor of *Arabidopsis thaliana* was performed along with the structure prediction (figure 3C and 3D). It was confirmed that 20 amino acid residues in REC domain are evolutionarily conserved. Among them, residues Asp15, Asp16 and Asp61 of MSMEG_0105 are involved in the phosphate binding.

The third family of LuxR/FixJ group of transcription regulators is the MalT-domain containing or LAL family (large ATP binding regulators of the LuxR family)^43^. This family of proteins includes proteins that contain an ATP-binding domain at the N-terminus and LuxR-type HTH at the C-terminus. This ATP-binding domain has conserved region called Walker A motif (GXXXXGKS/T, where X is any residue) that corresponds to P-loop ATPases^44^. Mtb has 1 MalT domain harboring protein (Rv0890c) and 3 ATPase domain containing proteins (Rv0386, Rv090c and Rv2488c). Rv0890c harbors both MalT and ATPase domains which could imply that ATPase is a part of MalT domain or it might harbor 2 ATPase domains. To confirm this, a sequence alignment for the three proteins with MalT protein of *E. coli* (used as a representative for LAL family of LuxR group) was performed (figure 3E). MalT of *E. coli* was considered as a representative for LAL family of LuxRs after sequence alignment for well characterized LAL domain harboring proteins (data not shown). All the three Mtb LuxRs harbor Walker A motif at their N terminus thus making all three, members of LAL family of LuxRs. Rv0890c which harbors both MalT and ATPase domains harbors only one Walker A motif indicating that both the domains (MalT and ATPase) are indeed a single domain but annotated differently. An early study on LAL proteins, included a set of 5 proteins in Mtb that harbor this domain^43^. These 5 proteins included the 3 ATPase domain containing LuxRs mentioned earlier and two new proteins (Rv0339c, and Rv1358). Surprisingly, these two proteins also harbor LuxR-type HTH thus expanding the list of LuxRs in Mtb. None of *M. smegmatis* LuxRs harbor MalT/ATPase domain.

The fourth and final family of LuxR/FixJ group of transcription regulators is the autonomous effector domain LuxR-type regulators^34^. This family includes proteins that do not have sensory domain but only harbors DNA-binding domain. MSMEG_1197 of *M. smegmatis* is only 85 amino acids in length consisting of HTH domain alone with no known annotated sensory domains and hence a member of this final LuxR family.

### LuxRs of mycobacteria harbor atypical ligand-binding domains

Mycobacteria also harbor domains which are usually not found in any of the LuxR/FixJ group of proteins. These domains include ATPase, AcyC, GAF, and PAS domains. Beginning with ATPase domain, which of certain proteins including LuxRs belongs to STAND (Signal Transduction ATPases with Numerous Domains) that are atypical AAA+ (ATPases associated with various cellular activities)^45^ of P-loop ATPases^46^ involved in signal transduction^47^. This type of ATPase is a part of LALs (explained in earlier section) and hence contains the consensus Walker A motif. Similar sequence alignment for the ATPase domain-harboring LuxRs (Rv0386, Rv0890c, and Rv2488c) was performed with RecA (a P-loop ATPase of *E. coli*) as shown in figure 4 which reconfirms that the 3 proteins indeed have ATPase domain of LALs. RecA of *E. coli* was considered as a representative for P-loop ATPases after sequence alignment for well characterized P-loop ATPase domain harboring proteins (data not shown).

**FIG 4.**
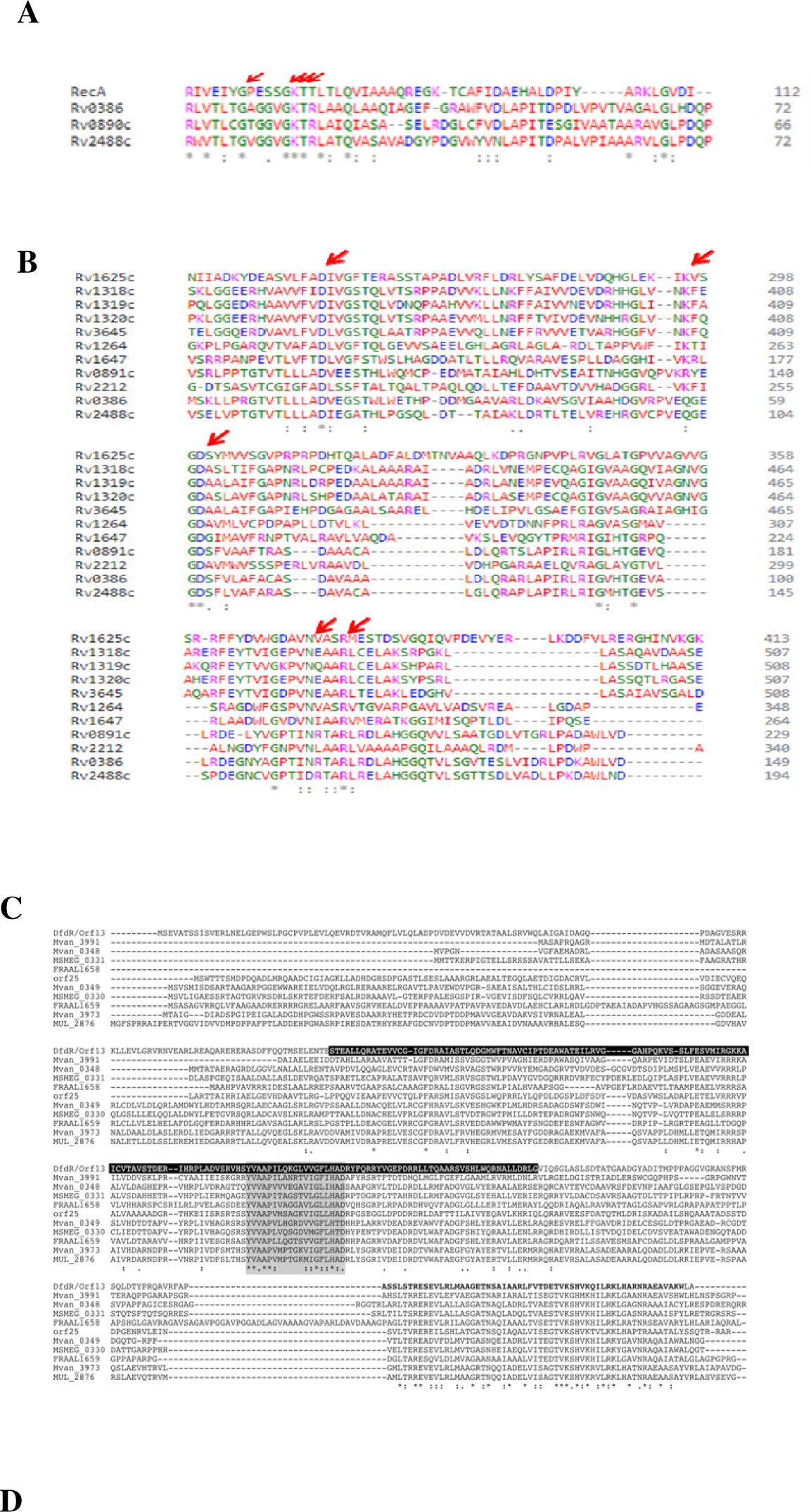
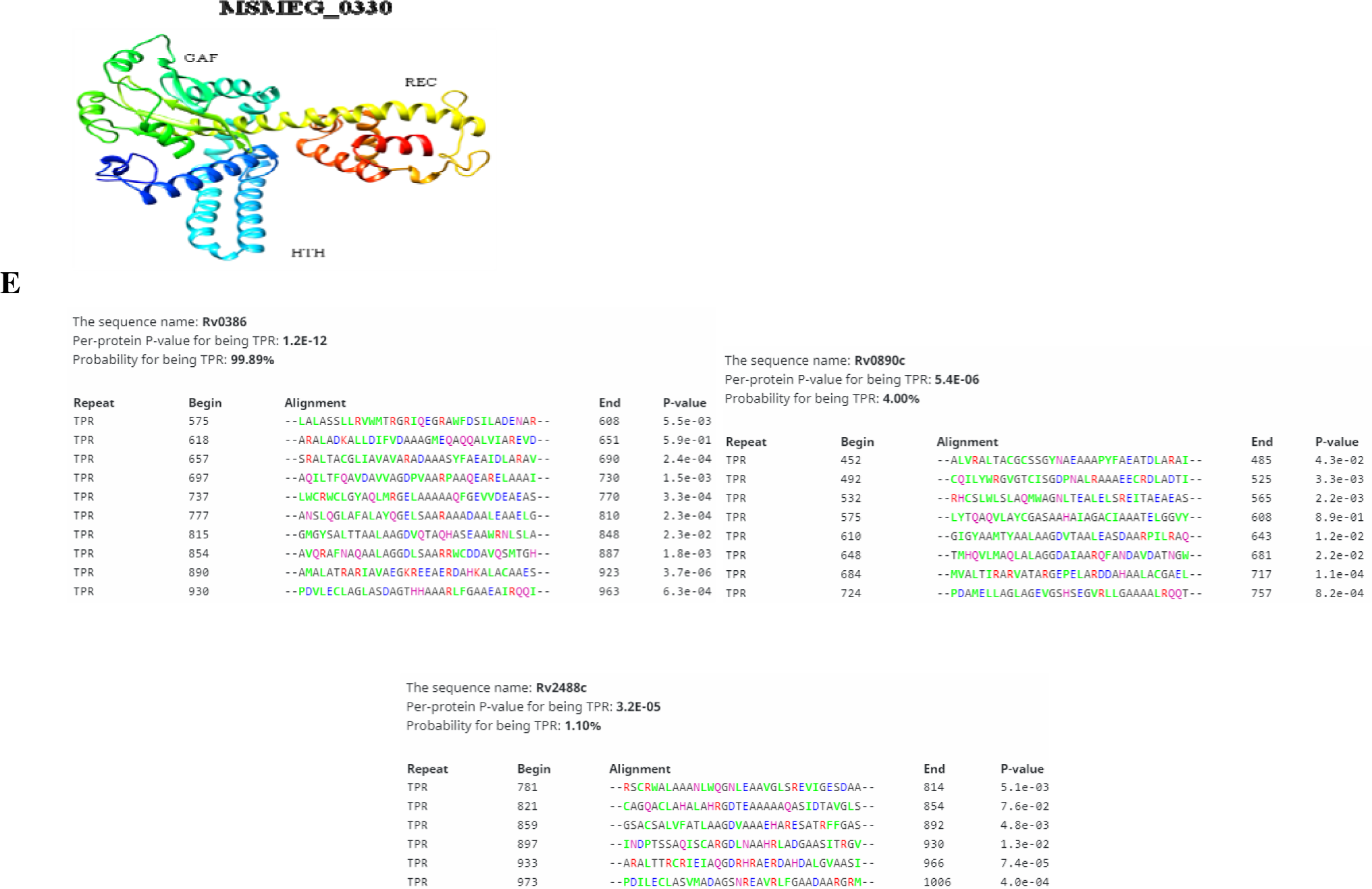
Multiple sequence alignment of atypical sensory domains of LuxRs in mycobacteria. Consensus amino acids are indicated by arrows. (A) *M. tuberculosis* has 3 LuxRs that harbor ATPase domain (Rv0386, Rv0890c, and Rv2488c). All the 3 proteins harbor a motif called Walker A motif (**G**XXXX**GKS/T**, where X is any residue) which is typical of P-loop ATPases, as that of RecA (a well characterized P-loop ATPase of *E. coli* used as a representative). (B) Sequence alignment of adenylyl cyclase (AC) domain-containing LuxRs (Rv0386 and Rv2488c) with other ACs of Mtb. Aspartate residues are involved in metal binding, aspargine and arginine residues are involved in catalytic transition-state stabilizing^50^. (C) Sequence alignment of MSMEG_0330 and MSMEG_0331 showing conserved amino acids in DNA binding domain. The amino acid regions highlighted in black background and bold type belong to a putative GAF domain and a DNA-binding domain of DfdR (transcription regulator), respectively (D) Predicted structures of MSMEG_0330 (used as a representative for GAF-harboring LuxRs) showing GAF domain along with REC and HTH motifs. Figures were drawn using Chimera^56^. (E) Prediction of TPR domain of Mtb LuxRs (Rv0386, Rv0890c, and Rv2488c) using TPRpred. Rv0386 with a probability of 99.89% seems to harbor TPR and hence might be involved in protein-protein interactions thereby regulating gene transcription. Rv0890c and Rv2488c have very less percentage of harboring TPR even being predicted to harbor the domain.

AcyC domain of LuxRs belongs to class III adenylyl cyclase (AC) which is known as universal (ancestral) class as they are also found in bacteria, archaea, and eukaryia^48^. Most of bacterial ACs belong to this category which is divided into subclasses IIIa to IIId^49^. Mtb has two LuxRs (Rv0386, and Rv2488c) that harbor AcyC domain and Rv0386 is a well-studied member of Mtb ATPase that belongs to class IIIc AC^50^. Sequence alignment for Rv2488c with other ACs of Mtb revealed the presence of consensus amino acids thus affirming the AcyC domain (figure 4B).

*M. smegmatis* has 3 LuxRs (MSMEG_0330, MSMEG_0331 and MSMEG_5651) that harbor an N-terminus GAF domain which is involved in ligand binding. GAF is an abbreviation for the first three characterized proteins that harbor this domain which include mammalian cGMP-binding phosphodiesterases, Adenylyl cyclases CyaB1 and CyaB2 from Anabaena sp., and *E. coli* Formate-hydrogen-lyase transcription activator^51^. Figure 4 (C) represents the REC and GAF domains of MSMEG_0330 and MSMEG_0331 with conserved regions for DNA binding^52^. MSMEG_0330 was used as a representative for GAF-harboring LuxRs for structure prediction as depicted in figure 4 (D). Sequence-structure alignments performed for these two sequences did not show any conserved amino acids within the active site or signal receiver domain. MSMEG_6568 of *M. smegmatis* harbors PAS domain, which is also a ligand binding domain similar to GAF. PAS domains were initially identified in and hence named after the first letter of these proteins which include circadian protein Period (per) and developmental regulator Sim (single-minded) of *Drosophila* and the vertebrate aryl hydrocarbon receptor nuclear transporter (ARNT)^53^. Interestingly, the PAS domains have a very low level of homology at the amino acid level where as their three-dimensional structure is highly conserved^54^. The sequence alignment with various PAS-domain harboring proteins did not reveal any consensus amino acids as expected.

Domain analysis for LuxRs using InterPro (as a confirmation for what has been predicted by CDD) revealed the presence tetratricopeptide repeat (TPR) motif in few LuxRs which was not predicted by CDD. TPR motif acts as interaction modules during formation of multi-protein complexes^55^. Rv0386, Rv0890c, and Rv2488c of Mtb harbor this domain. The presence of this domain was confirmed by using TPRpred which gives a P-value and the probability for harboring a TPR and the number of such repeats as shown in figure 4E. Only Rv0386 harbors TPR as predicted by TPRpred with the probability percentage of 99.89% and has 10 TPR units (figure 4E). Rv0890c and Rv2488c did not seem to harbor TPR as their probability percentage was less (4 and 1.1% respectively).

### LuxRs of mycobacteria harboring REC domain interact with histidine kinases

Few LuxRs harbor REC and TPR domains which are involved in phophotransfer and protein-protein interactions respectively. As mentioned in earlier sections, REC-containing LuxRs receive phosphate from a sensor kinase where in protein-protein interactions occur. These interactions with various proteins were predicted using STRING database. The interactive proteins for REC-harboring Mtb and *M. smegmatis* LuxR family proteins are shown in figure 5A and B respectively. Most of the proteins which are shown to be interacting with the REC-containing LuxR family proteins are HKs. Rv0386 of Mtb is the only LuxR that harbors TPR domain and hence the interactions of this protein were checked (figure 5C). Most of the protein that interact with Rv0386 are adenylyl cyclases (Rv1318c, Rv1319c, Rv1320c, Rv1647 etc.) indicating that TPR might form interacting platforms amongst various cyclases forming dimers or multimers culminating in transcription regulation.

**FIG 5.**
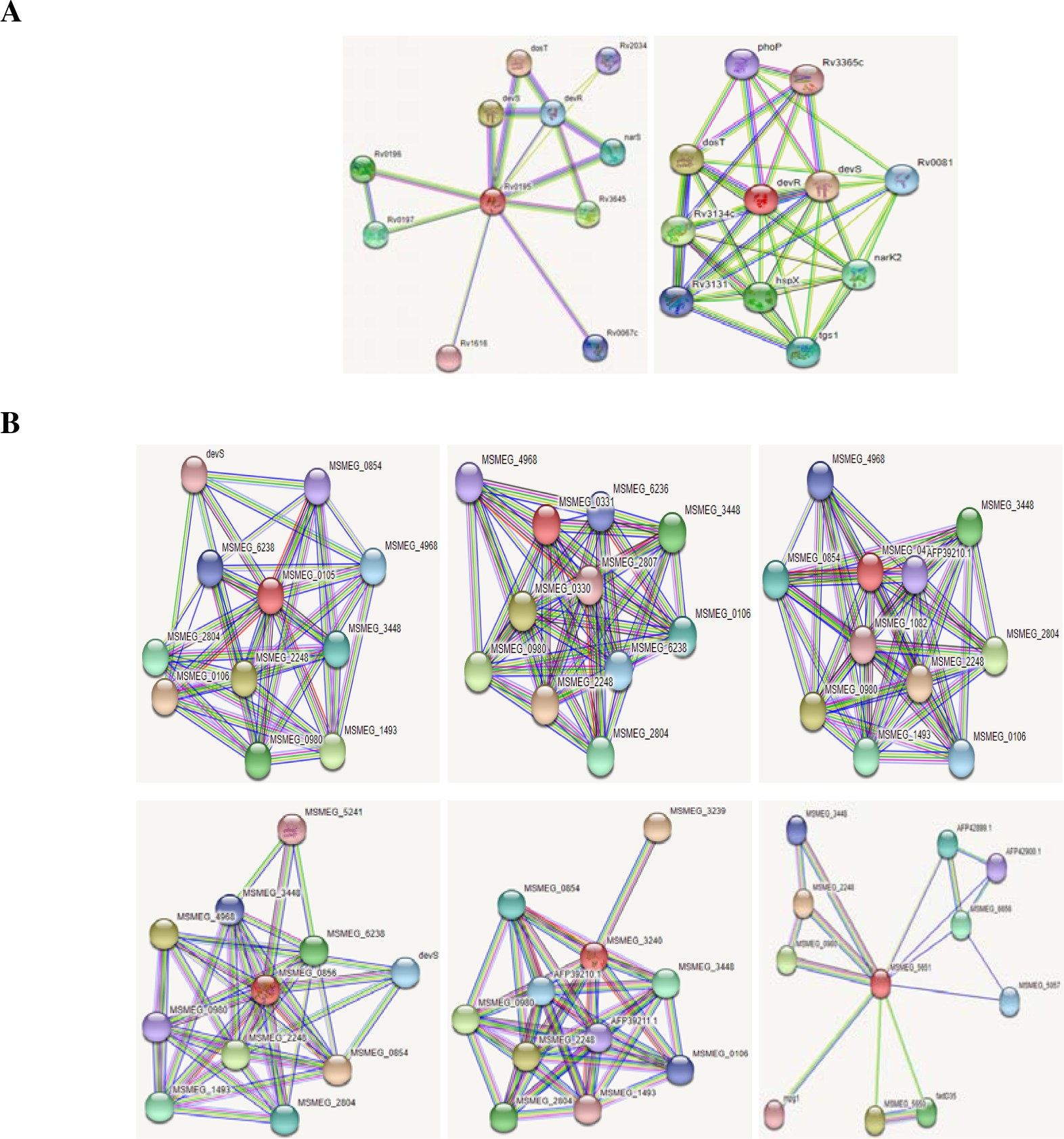
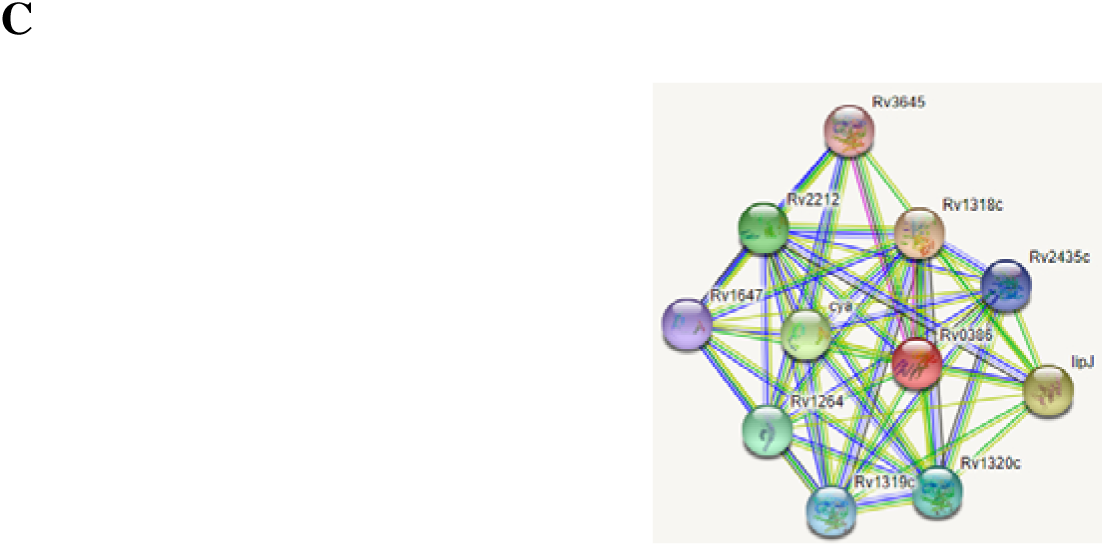
Interactions of the REC-containing LuxR annotated proteins of mycobacteria with other proteins using STRING for the identification of the histidine kinases (HK). Mtb (A) and *M. smegmatis’* (B) REC-harboring LuxRs interact with histidine kinases thereby accepting phosphate from HKs and thus modulating gene expression. (C) Interactions of Rv0386 (an adenylyl cyclase domain harboring LuxR of Mtb) with other adenylyl cyclases is probably mediated by TPR thus multimerize and regulate transcription. The query protein is indicated as red sphere in each of the interactome. The detailed list of interacting proteins with REC and TRP-harboring LuxRs is provided in supplementary table S6 and S7 respectively.

### GAF and PAS containing LuxR family proteins might interact with various ligands

Since three of the *M. smegmatis* LuxRs harbor GAF and PAS domains that bind to small molecules, the probable interactions of these proteins with different small molecules were predicted using STITCH online tool. MSMEG_0330, MSMEG_0331 and MSMEG_6568 didn’t seem to interact with any kind of small molecules whereas; MSMEG_5651 which harbors GAF domain was shown to probably bind to 3O-C12-HSL and c-di-GMP as shown in the figure 6.

**FIG 6.**
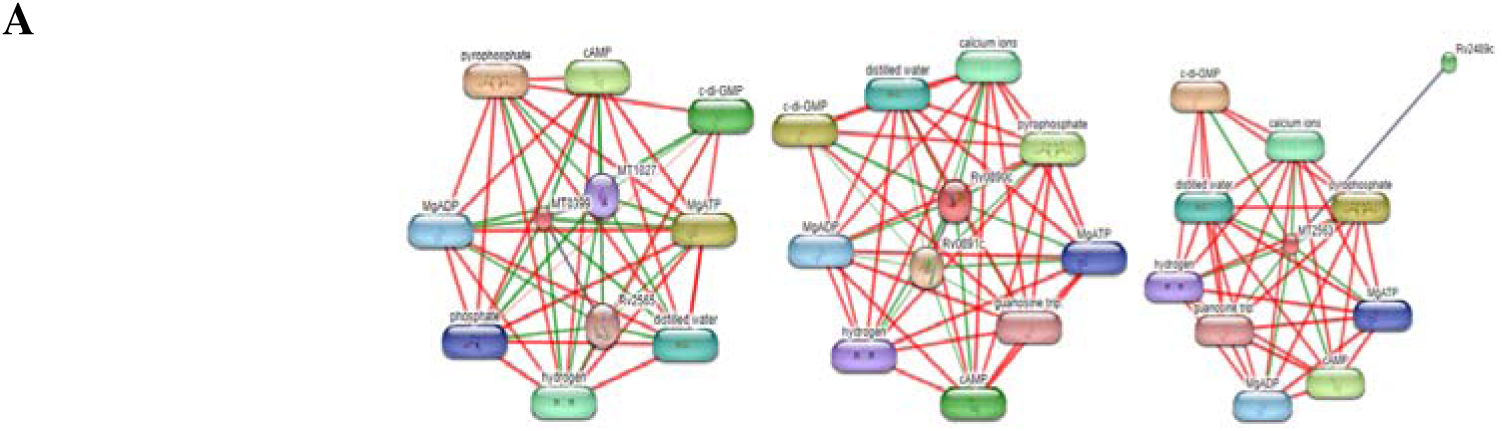
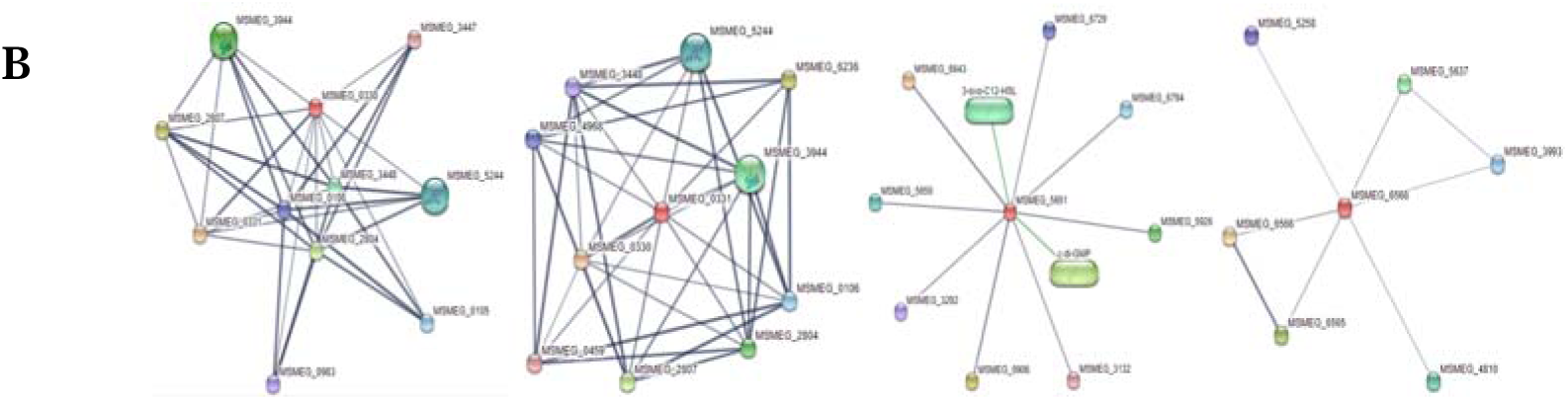
Predicted interactions of the ligand binding domain-containing LuxR annotated proteins of mycobacteria with various ligands. (A) *M. tuberculosis’* annotated LuxR family proteins Rv0386 (MT0399 in the figure), Rv0890c and Rv3133c (MT2563 in the figure) were predicted to interact with various small ligands like ATP, cAMP and c-di-GMP due to their harboring of cyclase domain and ATPase domains. (B) MSMEG_0330, MSMEG_0331 and MSMEG_6568 of *M. smegmatis* were shown not to interact with any of the small ligands where as MSMEG_5651 interacts with c-di-GMP and a homosrine lactone (3-oxo-C12-HSL). The latter molecule is one of the classical quorum sensing signals of *P. aeruginosa*. This hints for the probability of *M. smegmatis* detecting, and responding to AHLs from other bacteria. The query protein is indicated as red sphere in each of the interactome.

Apart from GAF, and PAS domains, few LuxRs which harbor cyclase and ATPase domains also interact with small molecules like ATP, cAMP, etc., as expected. Rv0386 and Rv2488c which harbor adenylate cyclase domain were shown to probably bind to MgATP, cyclic-di-GMP, and cAMP as shown in figure 6. The ATPase domain harboring LuxRs-Rv0386, Rv0890c, and Rv2488c were shown to interact with MgATP and MgADP as expected with ATPases.

### Mycobacteria harbor more proteins that have LuxR type HTH domain but have not been annotated as LuxR family of proteins

A further search in NCBI repository (based on the number of LuxR proteins in Mtb and *M. smegmatis* as listed by Santos CL, *et al*., 2012, from p2cs TCS components, LAL family of LuxRs^43^, and ACs in combination with HTH in Mtb^50^) revealed the presence of 3 more proteins in Mtb and 23 more proteins in *M. smegmatis* that have not been annotated as LuxR family proteins in the present annotation of Mycobacterial genome but harbor all the features of LuxR proteins. Their domain architecture is represented in supplementary figure S2. Orthologs of Mtb LuxRs in *M. smegmatis* and *vice-versa* have been looked for (data not shown) and found out that interestingly, MSMEG_3944, and MSMEG_5244 which were shown to be orthologs for Mtb annotated LuxR-Rv3133c, are indeed in this list of proteins that have not been annotated as LuxRs. Mtb protein Rv0844c has an ortholog in *M. smegmatis* which is MSMEG_0105 which is an annotated LuxR. Surprisingly, the ortholog of Rv0339c in *M. smegmatis*, MSMEG_0691 has the LuxR type HTH which earlier was not identified during the BLAST search in NCBI. The domain architecture of this protein is presented in supplementary figure S1. Thus, including this protein, *M. smegmatis* harbors 24 more proteins that have not been annotated as LuxRs. Considering the above analysis, very few LuxRs in Mtb have orthologs in *M. smegmatis* and *vice-versa* which might indicate that the proteins are not conserved within the genus and might have evolved in accordance to the adaptation of Mtb and *M. smegmatis* (pathogen and saprophyte respectively).

Most of these proteins harbor similar domains as that of annotated LuxRs like REC, MalT, ATPase, and AcyC in fusion with HTH. It is also of interest to note that few other proteins in this group harbor unique domains like AAA, AAA_16, TOMM, COG3899, HchA, and TPR. Similar domain analysis was carried out with this set of proteins as performed with the annotated LuxRs. Beginning with HTH domain, all the 3 proteins of Mtb and 24 of *M. smegmatis* have the typical LuxR/FixJ type HTH (data not shown) hence reaffirming their domain distribution and thus warranting their inclusion to the existing mycobacterial LuxR proteins group. So, combing all the proteins (annotated and that have not been annotated), Mtb has 8 and *M. smegmatis* has 33 LuxRs that have various sensory domains associated with HTH. Rv0844c of Mtb and 13 LuxRs of *M. smegmatis* were shown to harbor REC domain. Sequence alignment was performed (data not shown) which reveals that 3 proteins of *M. smegmatis* (MSMEG_1082, MSMEG_1301, and MSMEG_2807) do not harbor the consensus amino acids of the REC region and hence might not be a part of TCS. Mtb LuxR, Rv1358 and *M. smegmatis’* LuxRs (MSMEG_0321, MSMEG_1691, MSMEG_1901, MSMEG_2120, and MSMEG_4431) all harbor the Walker A motif of LAL family (data not shown) thus reassessing the presence of MalT domain. Surprisingly, domain analysis of Mtb Rv0339c using CDD revealed the presence of HTH domain only despite been categorized as LAL family^43^. Hence a sequence alignment with proteins belonging to LAL family was performed to confirm the presence of the domain. The analysis indeed revealed the presence of Walker A motif thus confirming its identity as LAL family of LuxR proteins. It’s ortholog in *M. smegmatis*; MSMEG_0691 also has similar domain architecture (LAL) (data not shown) thus making both of the proteins a part of LAL family.

AAA, AAA_16 and COG3899 are the ATPase domains and have the consensus region of P-loop ATPase as discussed earlier. Mtb has 1 LuxR and *M. smegmatis* has 8 LuxRs that harbor ATPase domain. All the proteins harbor the motif called Walker A motif (**G**XXXX**GKS/T**, where X is any residue) which is typical of P-loop ATPases, as that of RecA (a well characterized P-loop ATPase of *E. coli* used as a representative) (data not shown). The LuxRs with the ATPase domain hence might be an integral part of MalT/LAL family of LuxR group.

Rv1358 of Mtb harbors cyclase domain at its N-terminus which is confirmed by sequence alignment with characterized ACs of Mtb (data not shown). This protein along with annotated LuxRs, Rv0386, and Rv2488c harbor both cyclase and ATPase domains and hence might be involved in dual regulation of gene expression based on their catalytic action. TOMM system kinase/cyclase fusion protein domain containing proteins are fusions harboring a kinase domain and a cyclase domain followed by unknown domains. This domain is prominently present in members of *Burkholderia* and other bacteria and in genomic neighborhoods of a cyclodehydratase/docking scaffold fusion protein and a member of the thiazole/oxazole modified metabolite (TOMM) precursor family. MSMEG_0321, MSMEG_3290, MSMEG_4053, and MSMEG_6828 of *M. smegmatis* harbor this particular domain at the N-terminus. MSA of this set of mycobacterial proteins was performed with TOMM domain-harboring proteins (data not shown) and revealed the presence of consensus amino acids in this domain. The nature of this domain, its function in catalysis is completely unknown till date. MSMEG_4339 of *M. smegmatis* is the only LuxR predicted by CDD that harbors TPR domain but InterPro revealed that few other LuxRs (MSMEG_0321, MSMEG_1901, MSMEG_4205, and MSMEG_6828 of *M. smegmatis* and Rv1358 of Mtb) also harbor TPR. The confirmation for the domain was done using TPRpred which revealed that all the *M. smegmatis’* LuxRs (MSMEG_0321, MSMEG_1901, MSMEG_4205, MSMEG_4339, and MSMEG_6828) harbor TPR since their probability percentages were high (99-100%) (data not shown). Rv1358 of Mtb has very less probability percentage of 0.42% and hence might not harbor TPR. One peculiar atypical domain is “reg_near_HchA super family” domain in MSMEG_4053. This domain is only present in MSMEG_4053 at its C-terminus (supplementary figure S1 (B)). This domain is actually the DNA-binding domain (HTH) of LuxR family proteins that have chaperone/aminopeptidase HchA coding gene adjacent to the LuxR gene as a part of an operon^57^ and making it a new family within the LuxR/FixJ superfamily. In *M. smegmatis*, however the adjacent genes of MSMEG_4053 (MSMEG_4052, and MSMEG_4054) code for proteins that all are hypothetical of exceptionally smaller lengths (50 amino acids approximately) neither of which share similarity with HchA protein. Hence, MSMEG_4053 might not belong to this family but rather to the LAL family.

The interactions of REC and TPR-harboring Mtb and *M. smegmatis* LuxRs were checked using STRING database. Rv0844c (narL) - a *M. tuberculosis* protein was shown to interact with various proteins and most of which being HK- devS, narS, trcS, dosT and others (data not shown). *M. smegmatis’* proteins (MSMEG_0459, MSMEG_0983, MSMEG_1494, MSMEG_3447, MSMEG_3944, MSMEG_4378, MSMEG_4969, MSMEG_5244, MSMEG_5987, MSMEG_6236) interact with multiple other proteins most of them being histidine kinases (devS, MSMEG_0458, MSMEG_3448, MSMEG_2804, MSMEG_4968, 0980 and others). Hence, all the REC containing LuxRs probably act as response regulators of the two-component systems. MSMEG_0321, MSMEG_1901, MSMEG_4205, MSMEG_4339, and MSMEG_6828 harbor TPR domain and were shown to interact with different proteins, most of which being hypothetical proteins and very few being HK (data not shown). Interactions of the ligand binding domain-containing LuxRs of mycobacteria with various ligands were checked using STITCH. Rv0339c (MT0353) of Mtb was predicted to interact with ligands like 3-oxo-C12-HSL, and c-di-GMP although ATP was not included since the protein belongs to LAL family which harbors ATPase domain. Interactions of Rv1358 with different ligands including ATP, ADP, cAMP, and cGMP as the protein harbors both cyclase (AcyC) and ATPase domains (data not shown). *M. smegmatis*’ LuxRs (MSMEG_0321, MSMEG_0691, MSMEG_1691, MSMEG_1901, MSMEG_2120, MSMEG_3290, MSMEG_4053, MSMEG_4205, MSMEG_4431, and MSMEG_6828) containing ligand binding domains were shown to interact with different ligands that mostly are 3-oxo-C12-HSL, and c-di-GMP (data not shown). Both the molecules were shown to be quorum sensing signaling molecules in *P. aeruginosa* and *M. smegmatis* respectively. This might indicate a possible role of mycobacterial LuxRs as recognition system in inter-species quorum sensing.

## DISCUSSION

Quorum sensing enables bacteria to function as multi-cellular organisms by controlling the regulation of gene expression through QS signals which accumulate upon reaching certain cell density^58^. LuxR group of proteins are transcriptional regulators that mediate QS in gram negative bacteria by detecting and binding to AHLs which are the signaling molecules. LuxR proteins usually are found in association with a cognate protein-LuxI, which synthesizes AHLs. However, recent literature suggests that many bacterial species harbor “solo/orphan LuxRs” for which the cognate AHL synthases are absent. These LuxR solos are present in bacteria that synthesize AHL (for which there is a cognate LuxR to bind) but also in bacteria that do not synthesize AHLs^59^. These LuxR solos are known to detect and bind AHLs and non-AHL signals synthesized by the same cell or from different bacteria or host^60^.

LuxR and its homologs in various gram-negative bacteria belong to LuxR/FixJ superfamily proteins which contain a unique four helical HTH in their C-terminus. We employed computational approach to identify the domain architecture of LuxR family proteins in a clinically important pathogen *M. tuberculosis* and its saprophytic model-*M. smegmatis* and propose a new classification for few of the proteins that do not possess the characteristic HTH of LuxR/FixJ superfamily of transcription regulators. The proposed reclassification and their possible functional roles of LuxR family proteins in mycobacteria are illustrated in figure 7. LuxRs in mycobacteria could be classified broadly into four classes based upon the presence of different sensory domains that include 1) REC domain, 2) enzymatic domain, 3) ligand/protein binding domain, and 3) without sensory domains. A total of 7 proteins in Mtb and 10 in *M. smegmatis* have been annotated as LuxR family proteins, also a study had identified a total of 8 proteins in Mtb and 32 proteins in *M. smegmatis* that harbor LuxR type HTH DNA-binding motif^32^. Now, with a total of 10 proteins in Mtb and 34 in *M. smegmatis*, we have carried out the computational analysis for identification of the true nature of mycobacterial LuxRs. Combing all the proteins, 15 LuxRs (Rv0844c, Rv3133c, MSMEG_0105, MSMEG_0459, MSMEG_0856, MSMEG_0983, MSMEG_1494, MSMEG_3240, MSMEG_3447, MSMEG_3944, MSMEG_4378, MSMEG_4969, MSMEG_5244, MSMEG_ 5987, and MSMEG_6236) harbor REC domain that forms the first family, 15 LuxRs (Rv0339c, Rv0386, Rv0890c, Rv1358, Rv2488c, MSMEG_0321, MSMEG_0691, MSMEG_1691, MSMEG_1901, MSMEG_2120, MSMEG_3290, MSMEG_4053, MSMEG_4205, MSMEG_4431, and MSMEG_6828) harbor catalytic domain (ATPase and cyclase), 10 LuxRs (Rv0386, MSMEG_0321, MSMEG_0330, MSMEG_0331, MSMEG_1901, MSMEG_4205, MSMEG_4339, MSMEG_5651, MSMEG_6568, and MSMEG_6828) harbor ligand/protein interacting domain (PAS, GAF, and TPR), and 6 LuxRs (Rv0195, MSMEG_0473, MSMEG_1082, MSMEG_1197, MSMEG_1301, and MSMEG_2807) do not harbor any sensory domains. Interestingly, few LuxRs harbor multiple sensory domains which will be discussed later.

**FIG 7.**
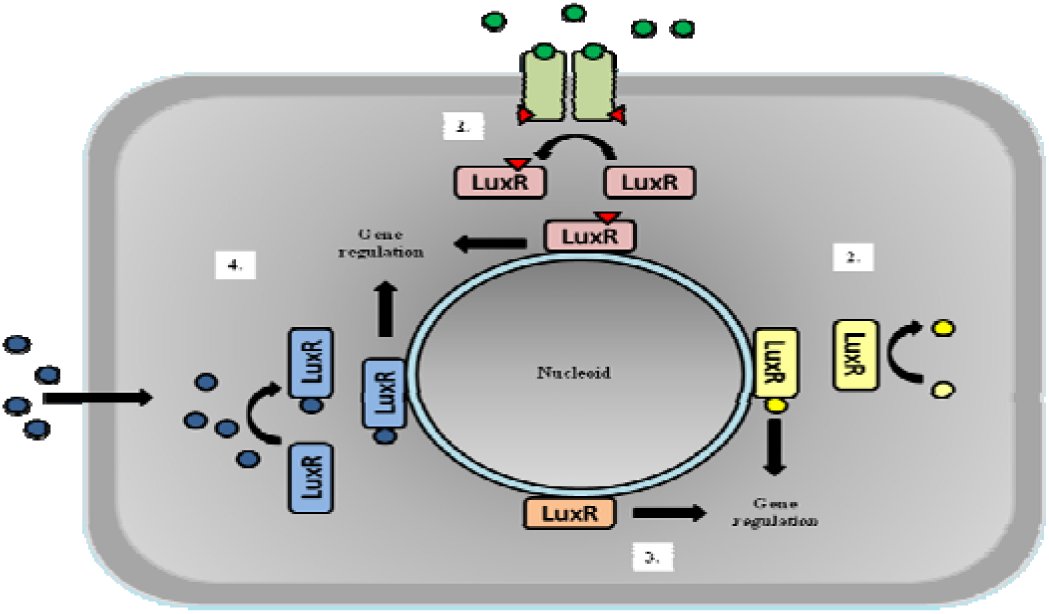
Representation of the mycobacterial LuxR superfamily proteins as different families based on their mode of activation (The LuxR in the figure indicates each LuxR family present in mycobacteria). (1) Some LuxR family proteins (Rv0844c, Rv3133c, MSMEG_0105, MSMEG_0459, MSMEG_0856, MSMEG_0983, MSMEG_1494, MSMEG_3240, MSMEG_3447, MSMEG_3944, MSMEG_4378, MSMEG_4969, MSMEG_5244, MSMEG_ 5987, and MSMEG_6236) function as response regulators of the two-component system activated by histidine kinases upon ligand binding. (2) Few proteins (Rv0339c, Rv0386, Rv0890c, Rv1358, Rv2488c, MSMEG_0321, MSMEG_0691, MSMEG_1691, MSMEG_1901, MSMEG_2120, MSMEG_3290, MSMEG_4053, MSMEG_4205, MSMEG_4431, and MSMEG_6828) harbor MalT domain with Walker A motif for ATPase activity. Included in this family is the other enzymatic domain AcyC– harboring LuxRs (Rv0386, Rv1358, and Rv2488c) that catalyze substrates hence regulating transcription. (3) The proteins harboring only HTH domain (Rv0195, MSMEG_0473, MSMEG_1082, MSMEG_1197, MSMEG_1301, and MSMEG_2807) without any of the sensory domains, also function as transcription regulators. (4) LuxRs harboring the small-molecule binding domains like PAS and GAF (MSMEG_0330, MSMEG_0331, MSMEG_5651, and MSMEG_6568) and LuxRs harboring TPR domain (Rv0386, MSMEG_0321, MSMEG_1901, MSMEG_4205, MSMEG_4339, and MSMEG_6828) that interact with different proteins, form the fourth family which binding to different targets (ligands and proteins), regulate transcription. Few LuxRs (Rv2488c, Rv1358, Rv0386, MSMEG_0321, MSMEG_1901, MSMEG_4205, and MSMEG_6828) harbor combinations of the different domains (AcyC+MalT, MalT+TPR, AcyC+MalT+TPR) thereby having multiple levels of regulation.

Out of 10 LuxRs in Mtb, 9 proteins harbor the HTH motif for DNA binding. Rv0894 gene product annotated as LuxR family protein product only has ATPase domain and no HTH motif. The protein is 393 amino acids in length and ATPase domain spans from 47 to 393 amino acids, almost the full-length protein. This indicates that the protein is not a transcription factor. There was no experimental evidence about the functionality of the protein either. Out of the remaining 9 proteins, RegX3 (Rv0491) though annotated as LuxR family and has an HTH motif, does not harbor the characteristic four helices of LuxR/FixJ superfamily but instead has winged HTH which is the characteristic of OmpR/PhoB family of RRs^61,62^. The wHTH is characterized by the presence of C-terminal β-strand hairpin which forms the wing and an open tri helix^61^. These wHTHs are present in several prokaryotic and eukaryotic transcription factors^63^. In Mtb, PhoP (Rv0757) which is a response regulator forming cognate pair with PhoR histidine kinase of TCS, harbors wHTH^64^ and is involved in Mtb virulence and division inside human bone-marrow derived macrophages^65^. RegX3 is also a response regulator of the two-component system with SenX3 (coded by Rv0490) being its cognate histidine kinase. The sensor kinase detects changes in phosphate levels thereby regulating transcription of several genes including the one coding for alkaline phsophatase^66^ and itself^67^ and the latter being the typical feature of AHL-binding LuxR proteins of gram negative bacteria. *M. smegmatis* which has 34 LuxRs on the other hand, has 33 proteins with HTH motif and one without the DNA binding motif. MSMEG_0545 which is of 175 amino acids only harbors the class 3 adenylate cyclase domain with no DNA-binding HTH motif. A study on the transcriptional regulation of MSMEG_0545 confirmed that the gene encodes for a functional protein. MSMEG_0545 was also shown to be up-regulated during starvation conditions and down regulated during H_2_O_2_ stress^68^. The first group of LuxR/FixJ superfamily of transcription regulation proteins includes the genes Rv0844c and Rv3133c of Mtb, and MSMEG_0105, MSMEG_0459, MSMEG_0473, MSMEG_0856, MSMEG_0983, MSMEG_1494, MSMEG_3240, MSMEG_3447, MSMEG_3944, MSMEG_4378, MSMEG_4969, MSMEG_5244, MSMEG_ 5987, and MSMEG_6236 of *M. smegmatis* harbor REC domains along with HTH DNA-binding motif. REC domain is phosphoacceptor domain of RRs that receives phosphate to an aspartate residue from a histidine residue in histidine kinase. Upon phosphotransfer, RRs undergo conformational changes and regulate signaling or catalytic activities thus making REC domain, a molecular switch controlling the response^69^. Rv3133c codes for DevR, which along with its cognate sensor DevS forms a two-component system that responds to different gaseous stresses including hypoxia^70^. DevR is activated upon phosphotransfer to aspartate from histidine either from DevS or DevT or from both forming a triad system that leads to the induction of various genes including DevR regulon^71^. From the known experimental evidence, it can be considered that this TCS whose RR being a LuxR homolog, might not be involved in QS but functions as a stress response system. Rv0844c is another REC-containing LuxR which is of 216 amino acids containing REC domain along with HTH. This has been annotated as NarL which is a known member of the TCS pair with Rv0845 (NarS) as its cognate HK. Similar protein in *E. coli* regulates gene expression in response to nitrate concentrations and also involved in nitrate metabolism under anaerobic conditions^72^. However, a study showed that in Mtb unlike *E. coli*, *narL* gene did not seem to affect the activity of nitrate reduction but had identified nearly 30 genes whose expression during aerobic growth was controlled by NarL in the presence of nitrate^73^. The same study also identified that DevR which is an important RR that synergistically regulates expression of itself and NarL during aerobic nitrate conditions and both the proteins interact with one another leading to a concerted gene regulation^73^. Taking the experimental evidence into consideration, NarL’s biological function sill remains elusive and the role of this TCS in QS would be a worthy question to be addressed. Another LuxR protein, MSMEG_0105 is of 216 amino acids which is the ortholog of Rv0844c. Till date, there is no experimental evidence of the functionality of this protein in *M. smegmatis* and remains to be studied. MSMEG_3944 annotated as DosR2 is an ortholog for Rv3133c (DevR) is of 217 amino acids. There are no studies available on this protein other than the expression of the gene was shown to be significantly repressed by PrrAB TCS in *M. smegmatis*^74^. This study involved the regulation of PrrAB TCS in M. smegmatis that represses certain genes involved in F_1_F_0_ ATPase and enhances expression of genes involved in respiration, hypoxia and iron regulation^74^. Although other than the expression profile for MSMEG_3944, there was no information regarding the functionality of MSMEG_3944 in *M. smegmatis*. The second ortholog of Rv3133c is *M. smegmatis* is MSMEG_5244 which is of 211 amino acids. The protein was shown to have a role in hypoxia response and mutation of this gene was shown to reduce the cell viability under hypoxia^75,76^. This protein was shown to activate hydrogen production thereby maintaining intracellular redox homeostasis^77^. The protein seems like a stress response regulative protein but whether it has a role to play in QS is unknown. These are no studies available for MSMEG_0459, MSMEG_0473, MSMEG_0856, MSMEG_0983, MSMEG_1494, MSMEG_3240, MSMEG_3447, MSMEG_4378, MSMEG_4969, MSMEG_5987, and MSMEG_6236 and hence the function of these proteins remains elusive.

The second group of LuxR/FixJ superfamily constitutes proteins which harbor enzymatic domains like cyclase and ATPase. The list consists of Rv0339c, Rv0386, Rv0890c, Rv1358, Rv2488c, MSMEG_0321, MSMEG_0691, MSMEG_1691, MSMEG_1901, MSMEG_2120, MSMEG_3290, MSMEG_4053, MSMEG_4205, MSMEG_4431, and MSMEG_6828 with ATPase domain (MalT) and Rv0386, Rv1358, and Rv2488c with cyclase domain. Rv0386 is 1085 amino acids long with both cyclase and ATPase domains. Rv2488c is one of the well studied LuxR of Mtb and belongs to class III adenylyl cyclases (AC)^50^. Interestingly it does not just have an AC activity but also has guanylyl cyclase activity which is dependent on the glutamine-asparagine pair^50^. Rv0386 was shown to be required for cAMP burst during infection resulting in production of TNF-α by macrophages^78^. Other than its cyclase activity, the ATPase activity of MalT domain was not studied and will be an active field to work upon. There are no functional studies available on Rv0339c, Rv0890c, Rv1358, Rv2488c, and none of *M. smegmatis’* LuxRs.

The third group comprises of proteins that do not possess any sensory domains and the classical example of this group is GerE of *B. subtilis*. GerE is the smallest member of LuxR/FixJ superfamily which is expressed during endospore formation of *B. subtilis*. This transcription factor either activates or represses Sig-K dependent transcription of several genes^33^. Rv0195 of Mtb and MSMEG_0473, MSMEG_1082, MSMEG_1197, MSMEG_1301, and MSMEG_2807 of M. smegmatis only harbor HTH at their C-terminus. Rv0195 is a well studied member which was shown to play a major role in Mtb dormancy and virulence^38^. This study was carried out with Mtb mutant for Rv0195 but the nature of the protein by itself as such was not studied which includes the different types of domains present and their function (except for the DNA-binding HTH domain), it’s mode of activation and action etc. Hence these aspects of the proteins still remain to be studied in detail to understand the role of such GerE-like proteins in dormancy and virulence. On the other side, the biological role for all the *M. smegmatis’* proteins remains unknown.

Many of the mycobacterial LuxRs harbor small molecule-binding and protein binding domains-GAF/PAS and TPR respectively. GAF and PAS are small-molecule binding domains. MSMEG_0330, MSMEG_0331, and MSMEG_5651 all harbor GAF domain at their N-terminus for which there is no literature available about their biological functions. MSMEG_6568 is the only mycobacterial LuxR that harbors a PAS domain. These are not experimental studies of the functionality of this protein. Surprisingly, the GAF/PAS domains share structural similarities autoinducer-binding domain of LuxRs that bind to AHLs^15,79^. The STITCH analysis for MSMEG_5651 also predicted interaction of this protein with 3O-C12-AHL. This might imply that at least few mycobacterial LuxRs bind and interact with AHLs although this has to be proved experimentally.

The protein-protein interacting domain TPR is found across domains of life. Rv0386 of Mtb and MSMEG_0321, MSMEG_1901, MSMEG_4205, MSMEG_4339, and MSMEG_6828 of *M. smegmatis* harbor TPR domain. Rv0386 not just harbors TPR but also possess cyclase and MalT/ATPase domains as discussed earlier in this section. Other than its role as cyclase, the TPR-dependent role of this LuxR in mycobacterial physiology is unknown. As with *M. smegmatis’*, TPR-harboring LuxRs, there is no literature available and hence their biological roles remain unknown. Interestingly, the TPR domain is also found in proteins that are involved in QS of gram-positive bacteria. These proteins belong to a group of QS receptors termed as RRNPP (Rap-Rgg-NprR-PrgX-PlcR) found in various gram positive bacteria^80^. These proteins are transcription regulators that bind to internalized peptide signaling molecules and regulate expression of QS genes involved in sporulation, biofilm formation, virulence etc^80^. The TPR region of these proteins is involved in peptide binding upon which these are conformation changes, thereby activating the proteins and enabling them to act as transcription activators by themselves or interacting with transcription regulators. ^80^. But, the HTH of these proteins is Cro/C1-type HTH which is present in the N-terminus of these transcription regulators consisting of 5 alpha helices and is very different from LuxR/FixJ type HTH that constitutes 4 helices^81^. Owing to the nature of TPR domain, its presence in mycobacterial LuxRs might indicate that these proteins either interact with other proteins or might bind to internalized messenger peptides as in QS of gram-positive bacteria.

From the literature and from our analysis it is very evident that indeed mycobacterial LuxRs are diverse with respect to sensory domains and hence their mode of activation. Owing to this diversity of sensory domains of mycobacterial proteins classified as LuxR family of proteins, and from the nature of such atypical domains, they are involved in various processes like dormancy and hypoxia and might be involved in other physiological functions. Although, there are many clues from the literature and from our analysis that they could possibly be involved in quorum sensing, yet the very phenomenon is underexplored in both *Mycobacterium tuberculosis* and *Mycobacterium smegmatis* with only information regarding biofilm formation in *M. smegmatis* being regulated by intra-cellular signals/second messengers. The nature of extracellular signals and the mode of action of the signal receivers like LuxRs needs to be studied in further detail to gain insights into the physiological aspects of LuxR group of transcription factors which help us to understand their importance in regulating bacterial gene expression either as a function of environmental/host induced stress or a quorum dependent mechanism of survival, symbiosis, and virulence.

## MATERIALS AND METHODS

### Sequence retrieval

Amino acid sequences for all 17 annotated (7 of *M. tuberculosis* and 10 of *M. smegmatis*) and 26 nonannotated (3 of *M. tuberculosis* and 23 of *M. smegmatis*) LuxRs were retrieved from UniProt^82^ (https://www.uniprot.org/) and Mycobrowser^83^ (https://mycobrowser.epfl.ch/) databases.

### Multiple sequence alignment

LuxR proteins of Mtb and *M. smegmatis* were aligned with classical LuxR family proteins for reconfirming the domain annotation by checking conserved amino acids, using the online tool Clustal Omega from EMBL-EBI (https://www.ebi.ac.uk/Tools/msa/clustalo/) web server.

### Domain analysis

Retrieved amino acid sequences for LuxR family proteins were investigated for the presence of functional domains using Conserved Domain Database (CDD)^84,85^ search. Also, the analysis was performed using InterPro^86^ (https://www.ebi.ac.uk/interpro/). Further, sequence-structure alignment was also performed using PROMALS3D tool which uses both the structural and sequence based constraints for aligning distantly related proteins^87^. This resulted in a consensus alignment of the target sequence with homologs with known structures. Both, the domain classification as well as the structural information were taken into consideration while performing alignment to obtain the structural and functional annotations. For the REC-harboring proteins (that are the response regulators of two-component systems), the confirmation for the REC domain and for identification of their cognate histidine kinases (if at all present (solo), paired or triad), the online server p2cs^88^ (http://www.p2cs.org/) was employed. For TPR domain prediction and confirmation, the online server TPRpred^89^ (https://toolkit.tuebingen.mpg.de/tools/tprpred) was employed

### Secondary structure prediction

The presence of helix-turn-helix and its nature was determined using SOPMA^90^ (https://npsa-prabi.ibcp.fr/cgi-bin/npsa_automat.pl?page=npsa_sopma.html)-an online prediction tool.

### Modeling 3D structure

To predict the 3D structure of the selected proteins (representatives for REC, and GAF-harboring LuxRs), comparative modeling algorithm RosettaCM was used which was accessible through the Robetta Server. This methodology uses an efficient conformational sampling technique along with the all-atom energy function which is physically realistic for better accuracy at atomic level^91^. A recent update of the algorithm enables the structure prediction through an integrated approach based on torsion space and Cartesian space template fragments. In addition, the loop modeling performed by an iterative fragment assembly and Cartesian space minimization, as well as high-resolution refinement are robust. In our study, MSMEG_0105 which belongs to the REC family had a close homology with signal receiver domain of the *Arabidopsis thaliana* ethylene receptor ETR1 (PDB ID: 1dcf) and hence we predicted the structure using comparative modeling. The sequences MSMEG_0330 and MSMEG_0331 belong to GAF and REC/GAF domains, respectively, were modeled using threading methods.

### Protein interaction prediction

Protein-protein interactions of REC–harboring, and protein-binding domains of LuxRs were determined using STRING database^92^ (https://string-db.org/). For predicting plausible ligands binding to ligand-binding domain harboring LuxR proteins (PAS, GAF, AcyC, MalT, ATPase, TOMM, AAA, AAA_16, and COG3899) online server STITCH^93^ (http://stitch.embl.de/) was used.

## SUPPLEMENTARY DATA

**FIG S1.**
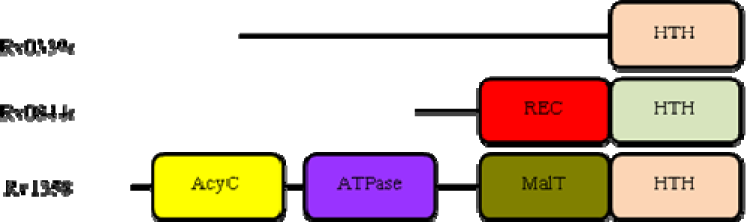
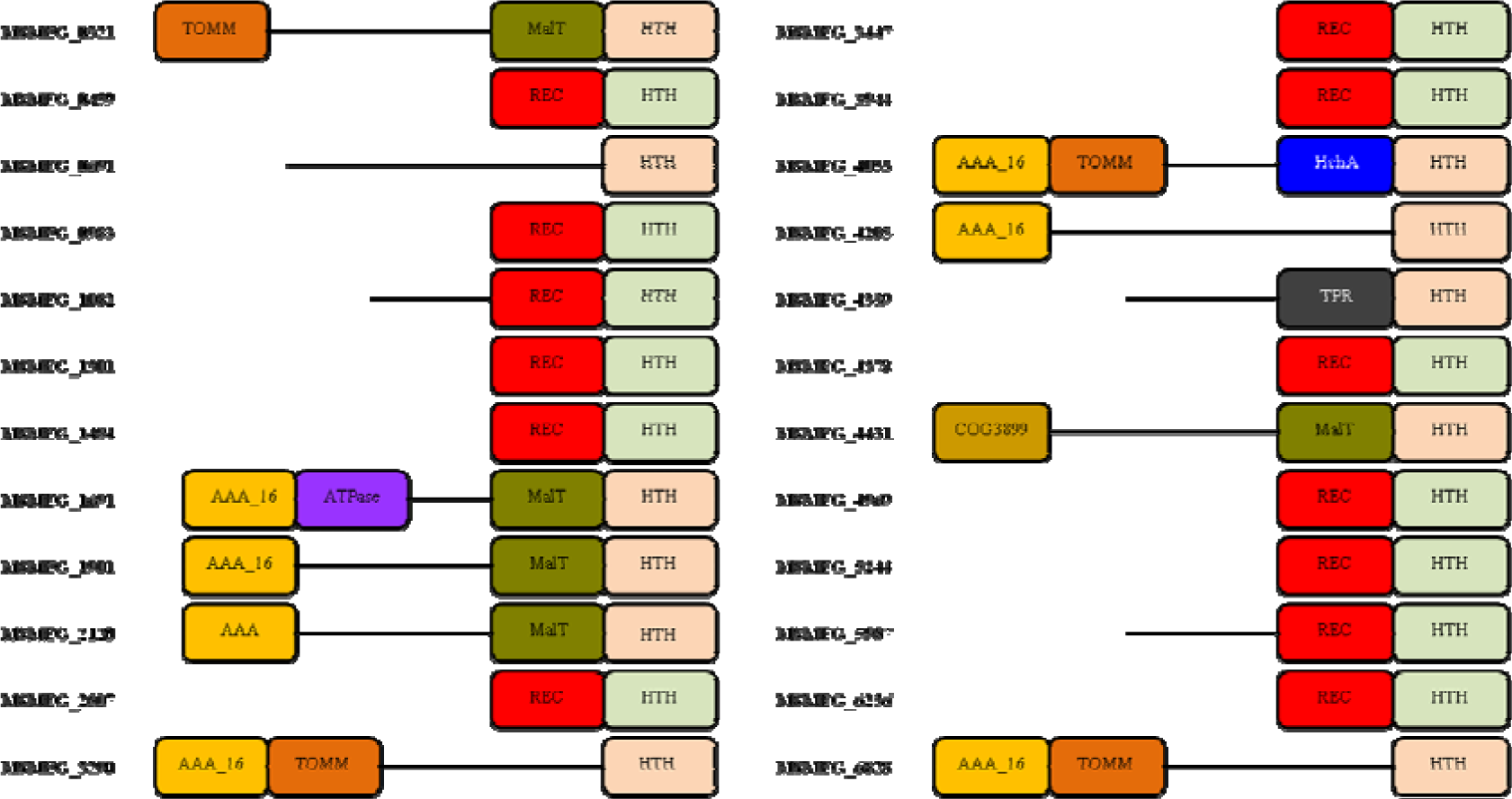
Domainal architecture of mycobacterial proteins that have not been annotated as LuxRs according to the present annotation in Mycobacterial proteome but harbor LuxR family DNA-binding domain. Representations of the domains and so the protein lengths are not to the scale. (A) *M. tuberculosis* has 3 proteins that are not annotated as LuxR family proteins. (B) *M. smegmatis* has 23 nonannotated LuxR family proteins. **HTH**-DNA binding helix-turn-helix motif, **REC** (receiver) or CheY-like phosphoacceptor is the signal receiver domain, **AcyC**-class 3 adenylate cyclase domain, **ATPase**-predicted ATPase domain, **MalT**-maltotriose, ATP binding domain, **TOMM-** TOMM_kin_cyc superfamily that contains a kinase, cyclase and an unnamed domain, **AAA_16**-AAA ATPase domain that contains a P-loop motif characteristic of AAA superfamily, **AAA**-ATPase (ATPases Associated with diverse cellular Activities), **HchA**-chaperone HchA-associated domain which is a protein deglycase, **TPR**-tetratrico peptide repeat region that mediates protein-protein interactions and the assembly of multiprotein complexes and **COG3899**-a predicted ATPase domain. The cartoon shows the diversity of domains associated with LuxR-type HTH and hints towards myriad of functions (based on functionality of the domain) that they might probably carry out.

